# Silver microwires from treating tooth decay with silver diamine fluoride

**DOI:** 10.1101/152199

**Authors:** Jong Seto, Jeremy A. Horst, Dilworth Y. Parkinson, John C. Frachella, Joseph L. DeRisi

## Abstract

Silver diamine fluoride (SDF) is a brush-on treatment for tooth decay that stops 81% of cavitated caries lesions (dental cavities). Before this innovation, caries was treatable only with operative approaches (dental fillings). SDF-treated lesions harden and become resistant to further decay. We hypothesized that the hardening is due to reaction with silver, rather than classic fluoride-mediated remineralization, because infected dentin is not amenable to remineralization. Using synchrotron microCT with 1.3 μm resolution, we observe filamentous densities up to 500 μm in length and 0.25-7.0 μm in diameter, formed *in situ* by applying SDF to caries lesions. We show that these “microwires” fill voids in the lesion caused by disease, and permeate through surrounding dentinal tubules. Using spectroscopy, we confirm that the chemical composition of the observed microwires is predominantly silver. To our knowledge, this represents the first structural microscale observations resulting from clinical SDF treatment. These novel observations hint at mechanistic explanations for the first clinical method to harden carious dentin besides remineralization. We hypothesize that SDF may not only achieve its antimicrobial functions by biochemical interactions, but also through its inherent ability to integrate into dentin.

## Introduction

Dental caries (tooth decay) occurs when plaque microbiota metabolize dietary sugars into acids that demineralize the tooth, exposing the collagenous dentin matrix, which is then degraded by endogenous and bacterial proteases (Featherstone, 2004). While unaffected or partially demineralized dentin forms a barrier, bacteria can invade further through the dentinal tubules. When enough bacterial toxins reach the pulp, they induce a painful vicious cycle of inflammation that results in necrosis, and through a permeable root, allowing the infection to ebb and flow with the serious consequences to spread along the facial fascial planes with systemic life-threatening consequences. Until recently, dental caries was only treated by excavation of the affected tissues and prosthetic replacement, via routine hard tissue surgery followed by restoration of form and function.

A non-invasive medical treatment for dental caries has been gaining rapid adoption by dentists. 38% w/v silver diamine fluoride (SDF) entered the United States market in 2015 as a medical device for treating tooth sensitivity. In 2016, an academic committee issued a recommendation for widespread off-label use of this topical solution to treat caries, based on 9 large long-term clinical trials (Horst et al., 2016). Later that year, application for an indication as a drug to treat caries received FDA Breakthrough Status, and in 2017 SDF was approved in Canada as an anti-caries agent.

Meta-analyses of published data sets show that simply drying a cavity then brushing on SDF arrests (stops) 81% of cavitated lesions (Gao et al., 2016). Due to simplicity and affordability, SDF is changing access to care for the most prevalent human disease. This is especially critical for treatment in young children, as the FDA recently released a warning that the sedatives commonly used to enable traditional operative dentistry (fillings) may have long term effects on cognitive development (Zhang et al., 2015).

It is thought that SDF arrests lesions via the silver ions lysing bacteria, and fluoride ions promoting remineralization to strengthen the lesion (Horst et al., 2016). However, fluoride has antibacterial action at high concentrations (Van Loveren, 2001), and *ex vivo* studies show more resistance to demineralization following treatment with SDF than with matching fluoride solutions without silver (Mei et al., 2013; Zhi et al., 2013). Thus, both silver and fluoride may contribute to both bactericidal effects and lesion strengthening.

SDF-arrested caries lesions are roughly twice as hard as normal dentin (Chu and Lo, 2008); this could not be caused by remineralization alone. Moreover, the outermost dentin is the most difficult to remineralize, due to matrix degradation. Reaction with the collagenous dentin matrix is supported by *in vitro* observations of silver particles on gelatin exposed to SDF (Lou et al., 2011).

Previous work on silver-based antibacterial compounds focused on silver nitrate (SN). While SN has been observed >1 mm deep in artificial lesions, dark SDF stains have been observed 200-300 μm into dentin (Shah et al., 2014). SN was used to treat dental caries until roughly 60 years ago, much as SDF is today. Studies on SN found semi-spherical polyhedral particles by light microscopy to course throughout the dentinal tubules in clinically treated artificial lesions (drill holes), penetrating as far as 1.65 mm, or 90% of the distance to the pulp (Zander and Burrill, 1943). SDF was developed to combine the anti-caries effectiveness of silver and fluoride, and outperformed each separately in clinical and laboratory models (Yamaga et al., 1972). Despite the positive anti-caries performance of SDF, a rigorous study of SDF penetration depth has been lacking. Approximately 45 years ago, exploratory microstructural work with scanning electron microscopy (SEM) of SDF-treated healthy dentin revealed sparse aggregates hundreds of microns into dentinal tubules (Shimooka, 1972). This study was not conducted on clinically treated teeth, and their findings of irregularly distributed particles were inconsistent with a structure that would spread mechanical loads to reinforce or harden lesions, as is observed clinically following SDF treatment. Thus, we investigated the deposition of solids in carious teeth following SDF treatment using advanced high energy synchrotron microcomputed tomography (microCT) and scanning electron microscopy with electron dispersive x-ray spectroscopy (SEM-EDS).

## Materials and Methods

### Collection of teeth

A convenience sample of 5 primary and 2 permanent teeth with natural cavitated caries lesions were collected following serial extraction for valid orthodontic purposes under UCSF BUA BU032969-03M. The dentist had treated by drying with air then scrubbing in SDF for 1 minute, from 1 month to 3 years later (13 months for the exemplar tooth) prior to extraction, whereupon they were kept in 0.1% thymol, sectioned to 2 mm thickness mesio-distally with a slow speed metallurgical saw, polished with sandpaper (Buehler, USA), and allowed to dry. Teeth were collected in accordance with de-identification procedures by the University of California, San Francisco Institutional Review Board.

### Optical light microscopy

Brightfield light microscopy was performed with an upright BX50 microscope (Olympus, Japan).

### SEM

Scanning electron micrographs were obtained using a Gemini Ultra-55 Analytical Field Emission SEM (Zeiss, Germany). An operating voltage of 10 keV was selected to differentiate the silver wires from dentin, whereas a lower voltage distinguishes peritubular from intertubular dentin (Supplemental Figure 1). Elemental analyses were conducted with an AzTec Energy Dispersive X-ray Spectroscopy system (Oxford Instruments, England) coupled to the SEM.

### Synchrotron Micro-Computed Tomographic Imaging

MicroCT was conducted at the Lawrence Berkeley National Laboratory Advanced Light Source (ALS) synchrotron at Beamline 8.3.2 (MacDowell et al., 2012). The ALS operates at 500mA ring current. An X-ray energy of 30 keV was selected using a multilayer monochromator (chosen to be above the 25 keV absorption edge of silver), with a 1 mm aluminum filter to prevent leakthrough of lower energies. Data was collected by imaging a Ce:LuAG scintillator with an optical Mitutoyo 5x lens onto a 14-bit scientific sCMOS camera with 2560x2160 6μm pixels (PCO, Germany), yielding 1.3 μm effective pixel size and a ~3.3 mm horizontal field of view. For each scan, a total of either 1025 or 2049 projections were acquired over 180° scan range. Samples were scanned separately with and without phase contrast for edge detection. Reconstruction was carried out using a custom ImageJ plugin for image preprocessing and Octopus Imaging Software (Inside Matters, Belgium) for tomographic reconstruction.

### Visualization of MicroCT Data

Reconstructed images were processed and compiled using Avizo 9.0 (FEI, USA).

## Results and Discussion

No silver or fluoride-related structures have explained the dramatic increase in lesion hardness following SDF treatment (Mei et al., 2014). In this study, light microscopy of sectioned SDF-treated primary teeth with natural caries lesions showed black stain where SDF absorbed and oxidized (Figure 1A), coating the dentin lesion outer surface and extending 700 μm pulpally, as elongated striations matching the orientation of the dentinal tubules. The pulpal extent of the stain follows a complex pattern, not simply filling all tubules (Figure 1A inset).

**Figure 1.**
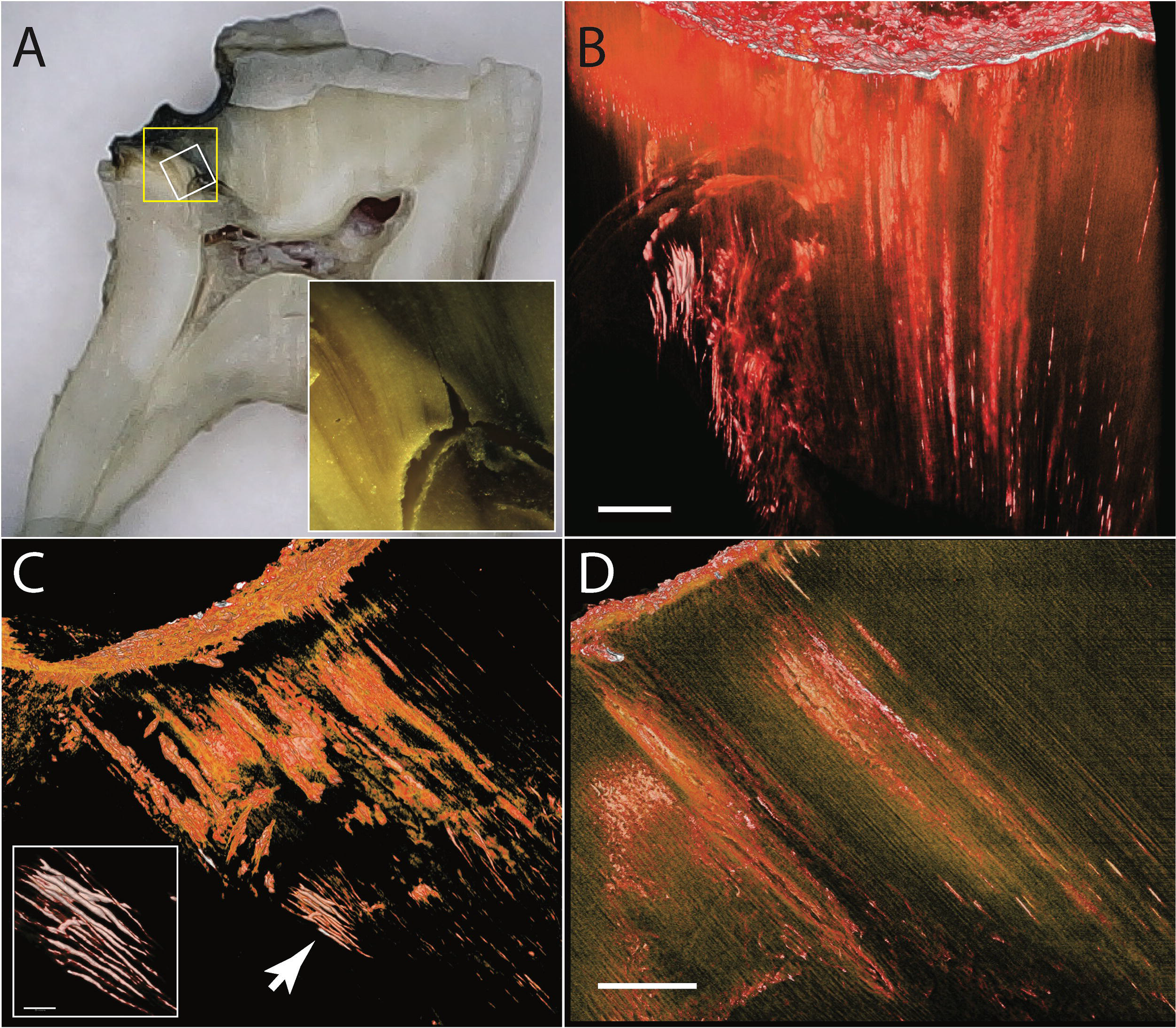
Microwires form in dentinal tubules following SDF treatment of caries lesions. A. Light imaging of primary tooth with natural caries lesion treated with SDF before extraction shows black and brown stains suggestive of silver deposition (5X, inset corresponds to white box: 30X). B. Synchrotron microCT imaging of the region of dentin outlined by the yellow box in panel A shows densities far greater than that of enamel, with a highly dense layer at the lesion surface, and structures of varying density extending pulpally. Volume rendering shown from occlusal, with visualization threshold set to show structures denser than dentin (100 μm scale bar). Colors in panels B-D denote increased density in order: yellow (density of enamel), orange, red, white. C. Shown from buccal as in panel A, with visualization threshold set to show structures denser than enamel. The distribution of silver matches that of the visible dark stains. Intensely dense wires form in the unaffected dentin (inset; 25 μm scale bar). Arrow points to region corresponding to inset. D. Scanning with phase contrast to enhance detection of solid-gas edges documents that the wires course along the dentinal tubules (100 μm scale bar). All images were taken of same area of same tooth section.

To characterize the shape and distribution of the structures observed as striated stains, we applied synchrotron microCT, which revealed densities far greater than that of enamel in every scanned tooth, matching the distribution and shape of stains observed by light microscopy. This manuscript describes analysis of one representative tooth; a comprehensive report on variations across all samples will follow. A highly dense <10 μm thick layer coats the surface (Figure 1B), with some discontinuities. Structures of varying density extend 700 μm pulpally from the surface, with most ending in cylindrical forms of intense uniform density (Figure 1B). Densities between that of enamel and dentin fills the first 175 microns (Figure 1B). Rendering to show only densities above that of enamel revealed irregular solids projecting generally in the orientation of dentinal tubules, and bridging across some tubules presumably broken down by the caries process (Figure 1C). Uniform filamentous densities of, roughly 100 μm in length and 3-7 μm in diameter, present near each other (Figure 1C inset) in structures reminiscent of bundled “microwires.” Less regular solids throughout the lesion range from 5-500 μm long. Rescanning using phase contrast confirmed that the densities do indeed follow the course of the dentinal tubules (Figure 1D).

These results document the penetration of SDF three times further into lesions than was previously thought (Shah et al., 2014). This is approximately the median of samples we have analyzed thus far. These structures were not observed despite previous attempts to image treated dentin using microCT, including studies demonstrating prevention of demineralization (Liu et al., 2012; Mei et al., 2013) or promotion of remineralization (Zhi et al., 2013), and evaluation of clinically treated successfully arrested teeth (Mei et al., 2014). The previous limitation of microCT appears to be the resolution; near-micron pixel sizes (1.3 μm) were necessary to image these structures. Indeed, teeth not treated with SDF imaged with the same microCT do not show these structures (e.g. Figure S1), nor does dentin distant from the treatment area in SDF-treated teeth (Figure S2 panel A). The shape and size of these microwires may enable distribution of forces across the dentin, reinforcing the lesion like rebar in cement. This observation has been made for peritubular dentin, increasing hardness and stiffness of dentin with its more dense form lining the dentinal tubules (Zaslansky et al., 2006; Weiner et al., 1999). However, the physical relationship of the wires to dentin substructures is not accessible at this resolution (Figures S2 panel B, S3 panel A).

SEM confirmed this novel observation of microwires, protruding from tubules near a tear in the dentin in a deep cross-section (Figure 2A), 250 nm in diameter - significantly smaller than observed using microCT. The differential electron density of silver enables application of high-energy x-ray and electron imaging techniques to probe the effect of SDF on treated tooth structures with submicron spatial resolution. Yet the observation of 3–7 μm diameter microwires by microCT, amidst detection of 250 nm microwires by SEM suggests that many thin microwires are below the detection limit of microCT with the settings used here.

**Figure 2.**
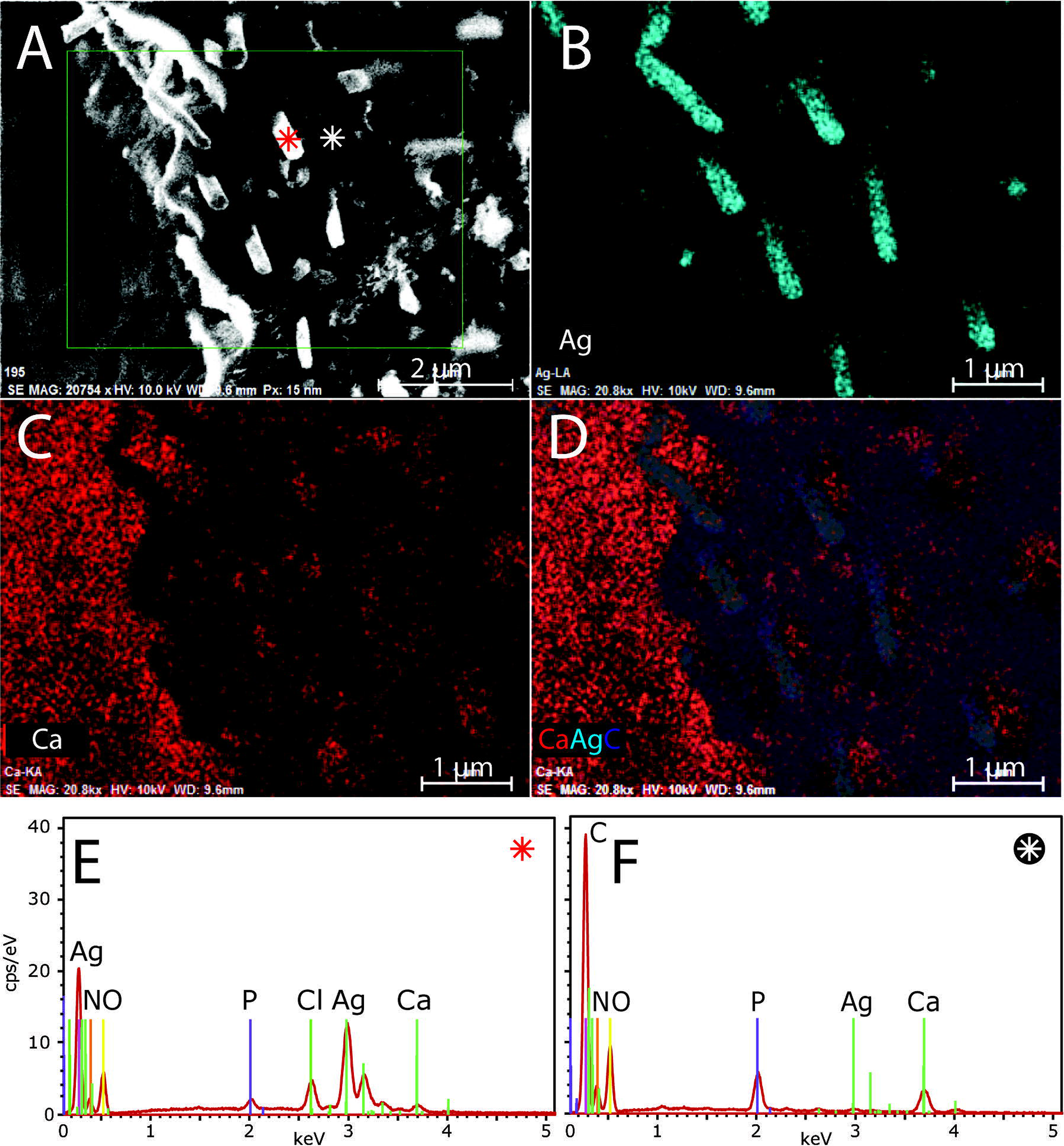
SDF-induced microwires are made of silver and chlorine. SEM-EDS imaging of a crosssection orthogonal to the dentinal tubules, showing A. the surface (electron density; 2 μm scale bar), B. silver, C. calcium, and D. overlay of silver (wires), calcium (to highlight peritubular dentin in longitudinal dimension), and carbon (to highlight intertubular dentin in cross-sectional dimension; 1 μm scale bars). EDS traces are shown for the positions corresponding to the E. red, and F. white asterisks in panel A. The microwires are comprised by approximately 4 parts silver to 1 part chlorine. Peritubular dentin is observed to the right of the wires in the tubules as high levels of calcium (red, panel C). *keV: kilo-electron volts, obs: observations, cps counts per second*.

While silver is the only element that can account for the densities observed in the microwires, the exact material composition remained unclear. Examination of the microwires with EDX spectroscopy revealed primarily silver composition (Figure 2B,E), and not calcium (Figure 2C), carbon, fluoride, oxygen, nitrogen, nor phosphorus (Figure S4). Overlay of silver, calcium, and carbon shows the wires descend into the tubules surrounded by highly calcified peritubular dentin (Figure 2D). Spectral characterization showed the microwires are comprised by silver and chlorine (Figure 2E,F), determined by ratiometric analysis to be approximately 4 parts to 1, respectively. While features of the dentin can be distinguished with SEM-EDS (Figure S3 panel B), more sensitive methods are required to determine the physical relationship of the wires to dentin substructures.

In summary, silver microwires 25-500 μm long and 0.25-7.0 μm in diameter are cast *in situ* in dentinal tubules by treatment of dental caries lesions with SDF. Diverse morphologies of the microwires are observed down to 700 μm from the surface, seeming to expand throughout dentinal tubules that were broken down by the caries process. The microwires may provide a reservoir of silver for antimicrobial action, prevent fluid flow through tubules, and increase hardness of the lesion. Hardness could arise from microwires distributing forces throughout the lesion and into the intact dentin. Fluid flow in tubules causes pain, and SDF clinically decreases sensitivity (Castillo et al., 2011).

SDF acts across several length-scales to treat dental caries lesions. At the level of the lesion, SDF hardens the lesion, perhaps by silver microwires forming in dentinal tubules, rather than by fluoride-driven remineralization. This is contrary to previous assumptions that fluoride strengthens the weakened tooth structure and silver merely acts as an antimicrobial agent.

Further work is needed to assess the correspondence of silver microwire physicochemical features to clinical outcomes. Structural characteristics such as microwire dimensions or distribution across tubules may be predictive of treatment success. Such measurements will be critical, given that there are presently no parameters to guide clinical protocols for maximizing effectiveness of treatment. Despite the need for further investigation, SDF is perhaps the most promising non-invasive intervention to control dental caries since the widespread implementation of water fluoridation.

## Acknowledgements

The Authors would like to thank Cate Quas and Steve Duffin for clinical samples; Howard Barnard for technical assistance, and Alireza Sadr for help with translation of references. This work was supported by NIH NIDCR grant T32-DE007306. The Authors declare no conflicts of interest related to the presented work.

